# Stimulus repetition induces a two-stage learning process in primary visual cortex

**DOI:** 10.1101/2024.09.03.611111

**Authors:** Lihan Cui, Ke Bo, Changhao Xiong, Andreas Keil, Mingzhou Ding

## Abstract

Repeated stimulus exposure alters the brain’s response to the stimulus. We investigated the underlying neural mechanisms by recording functional MRI data from human observers passively viewing 120 presentations of two Gabor patches (each Gabor repeating 60 times). We evaluated support for two prominent models of stimulus repetition, the fatigue model and the sharpening model. Our results uncovered a two-stage learning process in the primary visual cortex. In Stage 1, univariate BOLD activation in V1 decreased over the first twelve repetitions of the stimuli, replicating the well-known effect of repetition suppression. Applying MVPA decoding along with a moving window approach, we found that (1) the decoding accuracy between the two Gabors decreased from above-chance level (∼60% to ∼70%) at the beginning of the stage to chance level at the end of the stage (∼50%). This result, together with the accompanying weight map analysis, suggested that the learning dynamics in Stage 1 were consistent with the predictions of the fatigue model. In Stage 2, univariate BOLD activation for the remaining 48 repetitions of the two stimuli exhibited significant fluctuations but no systematic trend. The MVPA decoding accuracy between the two Gabor patches was at chance level initially and became progressively higher as stimulus repetition continued, rising above and staying above chance level starting at the ∼35th repetition. Thus, results from the second stage supported the notion that sustained and prolonged stimulus repetition prompts sharpened representations. Additional analyses addressed (1) whether the neural patterns within each learning stage remained stable and (2) whether new neural patterns were evoked in Stage 2 relative to Stage 1.

## Introduction

Repeated stimulus exposure alters the brain’s response to the stimulus (Sasaki et al. 2010). Behaviorally, repeating a stimulus, even over a short time period (e.g., a few dozen trials), can significantly improve performance in stimulus discrimination tasks (Fahle et al. 1995; Poggio et al. 1992). Similarly, in memory tasks, repeating a stimulus several times can lead to increased accuracy in memory recall (Oberauer et al. 2015). Over longer time periods, studies in both humans and animals have shown that after repeated exposure to visual gratings and objects for several days, the ability to discern grating orientations and object identities continues to improve (Jehee et al. 2012; Peissig et al. 2007). Understanding how learning and brain plasticity unfolds over different time horizons is an active area of neuroscience research.

Neurophysiologically, a repeating stimulus elicits decreased neural responses, a phenomenon known as repetition suppression. Repetition suppression as a robust form of perceptual learning (Henson 2003) has been observed in many brain regions and studied using a variety of measures in both humans and nonhuman primates (McMahon et al. 2007; De Baene et al. 2010; Demb et al. 1995; Buckner et al. 1995; Stern et al. 1996). In nonhuman primates, single unit responses to simple stimuli such as gratings decrease in the primary visual cortex after repeated stimulations (Kaliukhovich et al. 2018; Makino et al. 2015). In humans, fMRI repetition suppression, sometimes referred to as fMRI adaptation, has been observed in the primary visual cortex as well, especially when simple visual inputs such as contours and gratings are used as repeating stimuli (Kourtzi et al. 2005; Sapountzis et al. 2010; Fang et al. 2005; Larsson et al. 2016). These decreases in neural activity are believed to underlie improved perceptual efficiency through response adaptation to familiar stimuli (Henson et al. 2003; Adibi et al. 2013).

A number of neural models have been proposed to account for repetition suppression (Grill-Spector et al. 2006, Segaert et al. 2013). According to the fatigue model, repetition suppression occurs as the result of decrease in the firing rates of the stimulus-selective neurons (Miller et al. 1994; Müller et al. 1999; Westerberg et al. 2019), whereas the stimulus-non-selective neurons are less impacted (Grill-Spector et al. 2006), leading to decreased population-level activities. The sharpening model, in contrast, proposes that repetition prompts strong suppression of stimulus-non-selective neurons, with the stimulus-selective neurons being less impacted, resulting in decreased population-level activities but sharpened tuning for the repeated stimulus feature (Martens et al. 2012; Desimone 1996). This model has also been linked to the formation of increasingly sparse, efficient networks for representing the repeated feature (Gruber et al., 2002; Li and Keil. 2023). Both models, fatigue and sharpening, have been mainly tested with single-unit recordings in monkeys where stimulus selectivity can be precisely defined. In humans, the typical univariate fMRI analysis involves the averaging of BOLD activity across voxels in a brain region of interest, and as such, does not have the spatial resolution to test these models. A fMRI study performed at the single voxel level using complex visual objects as stimuli has provided evidence supporting the neural fatigue model in the ventral temporal cortex (Weiner et al. 2010). To what extent these models can be tested using fMRI in early visual cortex with simpler visual stimuli remains to be clarified.

The advent of multivoxel pattern analysis (MVPA) to fMRI data analysis offers a new perspective on testing the neural models of repetition suppression. Consider two repeating stimuli A and B (e.g., two Gabor patches of orthogonal orientations). Suppose that each stimulus is capable of evoking a distinct neural pattern in a cortical area (e.g., V1) and a SVM classifier can be trained to decode the two patterns. From the SVM classifier, one can derive not only the decoding accuracy indexing the difference between the A-pattern and the B-pattern but also a weight map, which divides the voxels into those that prefer A (higher BOLD activation for A than B) and those that prefer B (higher activation for B than A). Treating the A-preferring voxels and B-preferring voxels as “neurons” that are selective for A and for B, we can then examine the temporal dynamics of those voxels. In this MVPA-enabled analysis framework, the fatigue model predicts that the decoding accuracy between A and B will decrease with repetition, resulting from decreased activity in the stimulus-preferring voxels and thus diminished distinction between the two stimulus patterns, whereas the sharpening model predicts that the decoding accuracy between A and B will increase with repetition, resulting from decreased activity in the stimulus-non-preferring voxels and thus enhanced distinction between the two stimulus patterns. Our first objective in this work is to apply the MVPA approach to fMRI data to test the competing models of repetition suppression in the primary visual cortex.

The number of stimulus repetitions in typical repetition suppression studies is often small, on the order of 5 to 10 repetitions, after which the response typically becomes stabilized (Grill-Spector et al. 2001). One may interpret the finding to mean that learning is no longer taking place with additional repetitions of the stimuli. This interpretation, however, is at variance with extensive behavioral studies showing that perceptual abilities continue to improve over prolonged stimulus exposure, even in cases where the stimuli are not behaviorally relevant (e.g., the stimuli are being viewed passively). For instance, behavioral research has shown that after days of passive exposure to moving stimuli, individuals can exhibit improved ability to detect and discriminate coherent motion of stimuli (Watanabe et al. 2001; Watanabe et al. 2002). Neurophysiologically, research in nonhuman primates has shown that single unit activities in primary visual cortex continue to evolve with repeated presentations of the same stimuli over several days (Henschke et al. 2020). It is thus likely that the univariate fMRI approach lacks the ability to identify the learning processes taking place over time horizons beyond that typically employed in repetition suppressions studies. Our second objective in this work is to uncover these putative learning processes by applying the MVPA approach.

fMRI data were recorded from subjects passively viewing repeated presentations of two Gabor patches (45 degrees vs 135 degrees) in random order. Each Gabor patch was presented 60 times for a total of 120 trials. Both univariate and multivariate approaches were applied to analyze the data from the primary visual cortex. Particular emphasis was placed on (1) identifying the neural mechanisms of repetition suppression at the multivoxel pattern level and (2) exploring learning processes beyond the time scale used in typical repetition suppression studies.

## Materials and Methods

### Participants

The experimental protocol was approved by the Institutional Review Board (IRB) of the University of Florida. Eighteen healthy individuals (nine females and nine males; mean age: 21.3±4.5) gave written informed consent and participated in the study.

### Experimental paradigm

The stimuli were presented on a back-illuminated screen (60 cm × 60 cm) placed 230 cm away from the participant’s head. The participant viewed the stimuli through a set of prismatic glasses attached to the radio frequency head coil. Two Gabor patches with orthogonal orientations (45° and 135°), referred to as Gabor 1 (45°) and Gabor 2 (135°), were used as the stimuli. In each trial, the stimulus was displayed for 1s, which was followed by a randomized inter-trial interval (3, 5 or 7s) (Figure 1A). The subject was asked to passively view the stimulus. There were a total of 120 trials, 60 trials for each of the two Gabor patches; the order of stimulus presentation was randomized.

**Figure 1.**
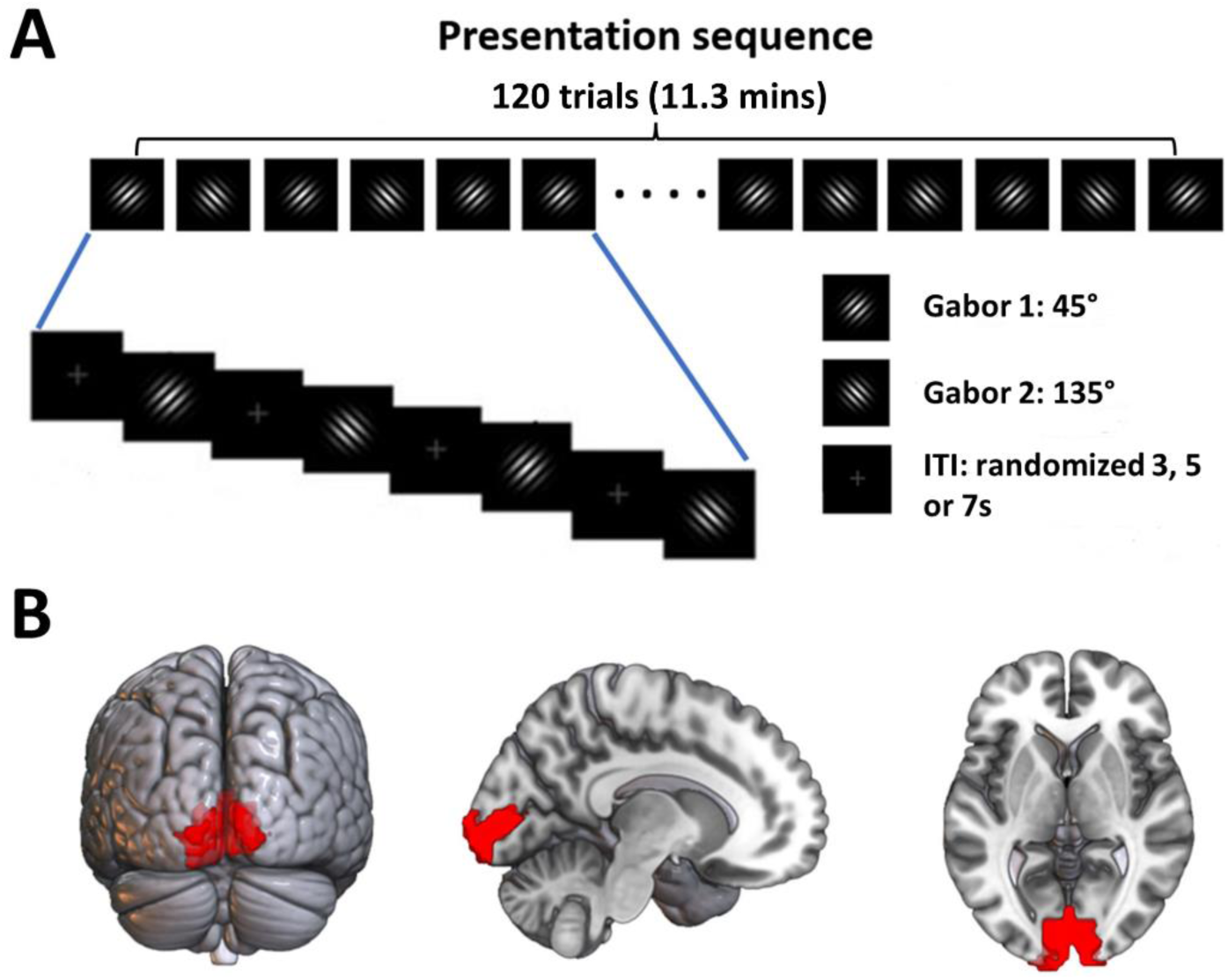
Experimental paradigm and ROI definition. **(A)** Timeline of stimulus presentation. There were two Gabor patches, Gabor 1 (45°) and Gabor 2 (135°), and each was presented 60 times for a total of 120 trials. The order of stimulus presentation was randomized. **(B)** Primary visual cortex (V1), defined according to a retinotopic atlas of the visual cortex, was chosen to be the region of interest (ROI). ITI: Inter-trial interval.

We note that the data analyzed here came from the habituation session of a Pavlovian fear conditioning paradigm consisting of three sessions: a habituation session (the present data), an acquisition session in which the CS+ (Gabor 1) was occasionally paired with a human scream (the unconditioned stimulus or US), and an extinction session in which 120 trials of the two Gabor patches were presented in random order without the US. Two manuscripts have been published on these data to investigate the neural mechanisms of fear learning and extinction (Yin et al. 2020; Yin et al. 2018). The questions addressed here have not been addressed in those previous studies.

### Data acquisition

The fMRI data were acquired on a 3-Tesla Philips Achieva MRI scanner (Philips Medical Systems, Netherlands) with the following parameters: TR of 1.98s, TE of 30ms, flip angle of 80, field of view of 224mm, 36 slices, and 3.5 × 3.5 × 3.5mm voxels. A T1- weighted structural image was also obtained from each participant.

### Data preprocessing

The fMRI data were preprocessed with SPM. Slice timing correction was first done using the interpolation method. The images were spatially realigned to the first stable volume, normalized and registered to the Montreal Neurological Institute (MNI) template, and resampled to a spatial resolution of 3mm by 3mm by 3mm. The transformed images were spatially smoothed with a 7 mm full width at half maximum Gaussian kernel. The BOLD time series were high-pass filtered with a cutoff frequency at 1/128 Hz to remove low frequency temporal drifts.

### Region of interest

The primary visual cortex (V1) was chosen as the region of interest (ROI). The V1 ventral and V1 dorsal of both hemispheres from a probabilistic visual retinotopic template (Wang et al. 2015) were combined to form the V1 ROI (see Figure 1B).

### Single-trial estimation of BOLD response

The single-trial BOLD activation was estimated by the beta series method based on the general linear model approach to event-related fMRI data analysis (Rissman et al. 2004). Specifically, in the general linear model, each trial was represented by a separate regressor, and six head motion regressors were included as regressors of no interest to account for potential movement artifacts. 120 single-trial level BOLD activation patterns within the ROI enabled the trial-by-trial analysis of neural activity change with learning.

### Univariate analysis of fMRI data

For each subject, BOLD activation within V1 for each trial was averaged across all V1 voxels, which were then further averaged across eighteen subjects to generate trial-by-trial population level BOLD activation for V1. Systematic changes in univariate activation over trials were examined for evidence of repetition suppression and/or possibly other forms of learning.

### MVPA analysis of fMRI data

*Moving window approach:* The differences in multivoxel responses to Gabor 1 and Gabor 2 were examined using MVPA applied at the subject level. To study the temporal dynamics of multivoxel representations, we adopted a moving window approach, in which a time period of interest (e.g., the time period of repetition suppression) was divided into overlapping analysis windows with each window consisting of an equal number of consecutive Gabor 1 and Gabor 2 trials and two neighboring windows separated by one trial (i.e., stepping the analysis window forward one trial at a time). For example, as shown in Figure 2A, the time period during which repetition suppression was observed was 24 trials in duration (12 repetitions each for Gabor 1 and Gabor 2). The first analysis window consisted of Gabor 1 repetitions 1 to 7 (G1R1 to G1R7) and Gabor 2 repetitions 1 to 7 (G2R1 to G2R7), the second analysis window consisted of Gabor 1 and 2 repetitions R2 to R8, …, and the sixth analysis window consisted of Gabor 1 and 2 repetitions R6 to R12. Within each analysis window, the support vector machine method as implemented in the LibSVM package (Chang et al. 2011) was combined with cross-validation to yield a decoding accuracy for classifying Gabor 1 vs Gabor 2 based on the patterns evoked by the two stimuli. As the analysis window was stepped forward, the above process was repeated for the new analysis window, yielding another decoding accuracy. Across all windows a decoding accuracy time course was obtained for the subject. The population level decoding accuracy time course was obtained by averaging the subject level decoding accuracy time course across subjects and taken to indicate learning-related changes in multivariate representations of the Gabors.

**Figure 2.**
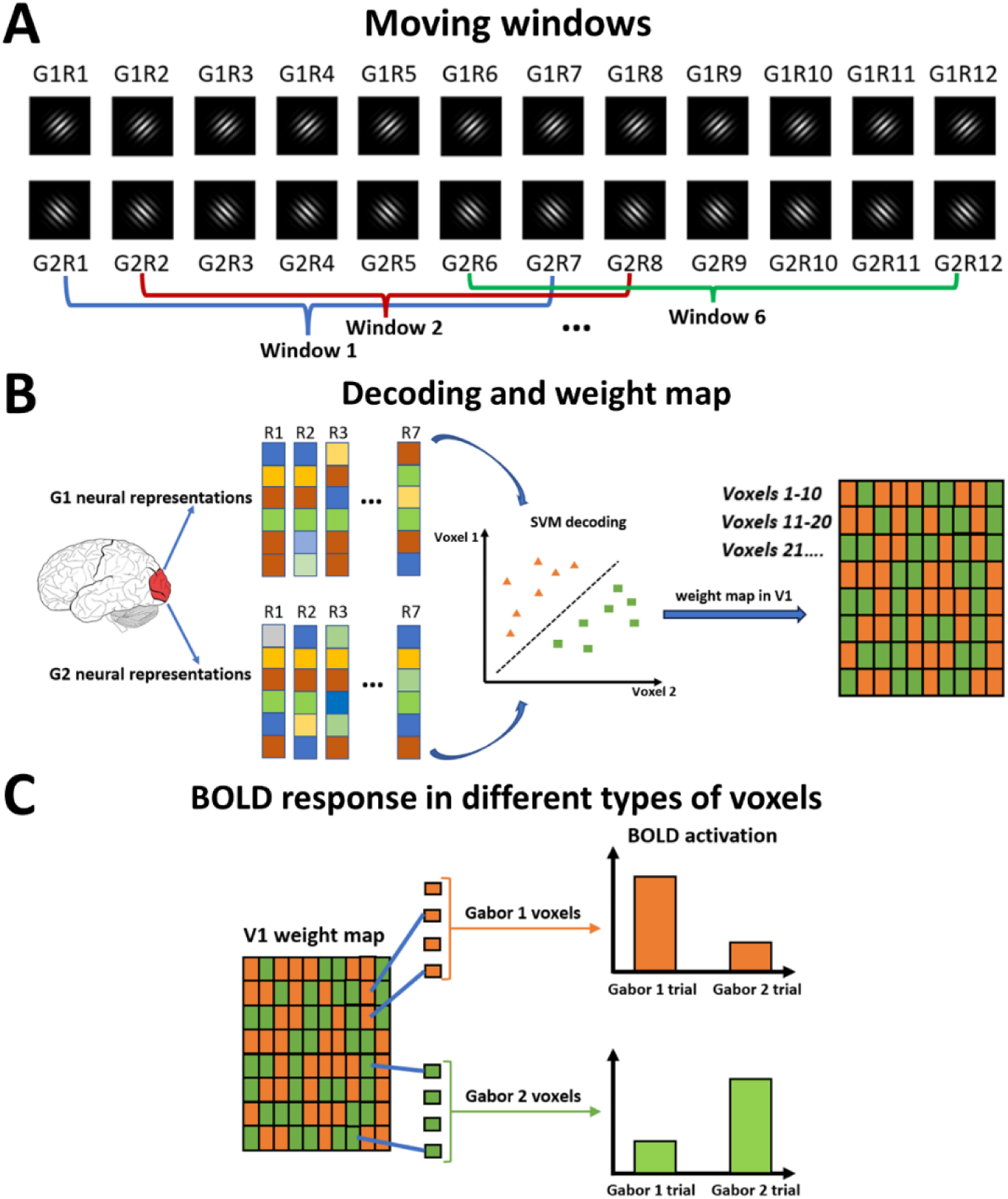
Moving-window MVPA analysis. **(A)** Definition of moving windows (G1R1 indicates first presentation of Gabor 1, G2R1 first presentation of Gabor 2, and so on). **(B)** SVM classification and weight map within a window. In addition to decoding accuracy, the SVM classifier also generates a weight map, which defines two types of voxels: Gabor 1 voxels (orange) and Gabor 2 voxels (green). **(C)** Gabor 1 voxels and Gabor 2 voxels are selective for Gabor 1 and Gabor 2 respectively.

*Statistical assessment of decoding accuracy:* The Wilcoxon sign-rank test was utilized to assess the statistical significance of the decoding accuracy. Specifically, the decoding accuracy from each analysis window was compared against a chance level of 50%, and the resulting p-values were corrected for multiple comparisons using a false discovery rate (FDR) control threshold of p < 0.05. Cluster size (>=3 points) was further imposed to enhance statistical rigor.

### Weight map analysis

*Voxel level stimulus selectivity:* In addition to detecting the differences in stimulus-evoked patterns, the MVPA also makes possible the determination of stimulus selectivity at voxel level, opening the possibility for testing the models of repetition suppression using fMRI. For a SVM classifier, the weight vector is the vector that is normal to the hyperplane that maximally separates the data points of two different classes (e.g., Gabor 1 and Gabor 2). This vector has the same dimension as the number of voxels in a ROI (e.g., V1). The physiological interpretation of the classifier weight vector is not straightforward. Voxels that do not contain task-related signals can be given high weight whereas voxels that contain task-related signals may be given low weight. Haufe et al (Haufe et al. 2014) proposed a transformation that transforms the weight vector from the SVM classifier into a new vector according to, A = Covariance(X) * W, where the X is the recorded data, W is the weight vector from the SVM classifier, and A represents the transformed weight vector. The new weight vector, referred to as the weight map here, is shown to reflect more accurately the generative neurophysiological processes behind stimulus patterns (Haufe et al. 2014; Grootswagers et al. 2017). For the purpose of this work, we used the weight map to identify stimulus selectivity at the voxel level; see Figure 2B. Specifically, the sign of the weight in each voxel obtained from the weight map provides information about which of the two stimuli (Gabor 1 vs Gabor 2) the voxel is selective for, and the magnitude of the weight denotes the strength of the selectivity. To illustrate, suppose that the Gabor 1 is assigned the sign of +1 and the Gabor 2 the sign -1. A voxel with a positive weight indicates that the voxel has higher activation for Gabor 1 compared to Gabor 2 and is thus deemed to be selective for Gabor 1. A voxel with a negative weight will have an opposite activation profile and is thus deemed to be selective for Gabor 2. See Figure 2C for an illustration. In both cases the magnitude of the weight determines the magnitude of differential activation.

### MVPA formulation of repetition suppression models

*Temporal dynamics of decoding accuracy:* The fatigue and sharpening models make distinct predictions regarding the dynamics of the decoding accuracy between the two Gabors. According to the fatigue model, the decoding accuracy between the two repeating Gabors should decline as learning progresses, as stimulus-selective neurons reduce their response amplitude and the activity of stimulus-non-selective neurons remains relatively stable, leading to less distinctively different neural patterns between the two stimuli; see Figure 3A (top). In contrast, according to the sharpening model, the decoding accuracy should increase with increasing stimulus repetition, due to the fact that the stimulus-non-selective neurons decrease in activity with learning and the activity of stimulus-selective neurons stays relatively stable, leading to more distinctively different patterns between the two stimuli; see Figure 3A (bottom).

**Figure 3.**
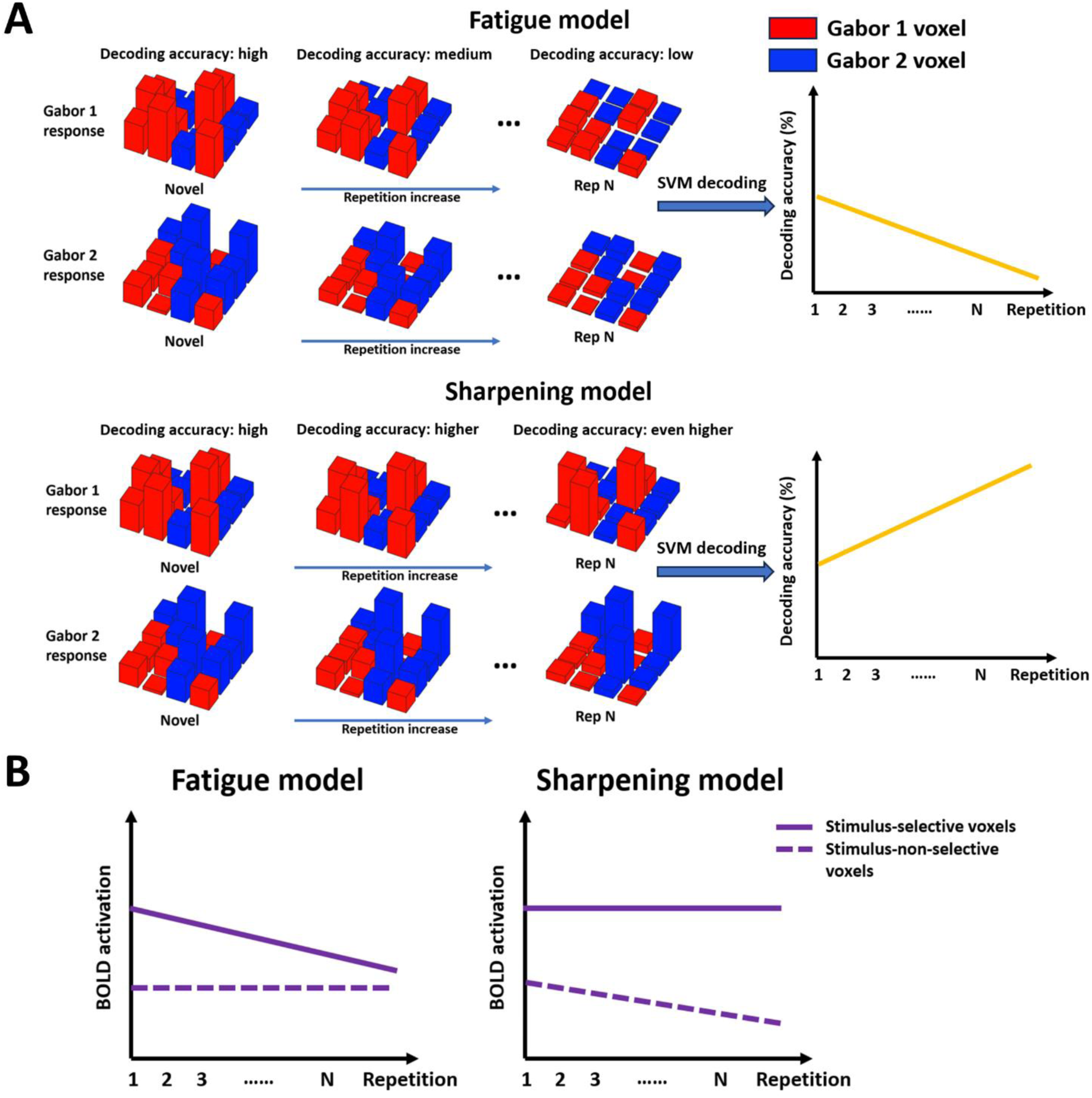
MVPA formulation of fatigue and sharpening models. **(A)** The fatigue model predicts that the two patterns evoked by Gabor 1 and Gabor 2 become less distinct with stimulus repetition, resulting in decreased decoding accuracy (top). The sharpening model, in contrast, predicts that two patterns evoked by Gabor 1 and Gabor 2 become more distinct with stimulus repetition, resulting in increased decoding accuracy (bottom). **(B)** The fatigue model predicts that BOLD activity in stimulus-selective (stimulus-non-selective) voxels decreases (stays unchanged) with stimulus repetition (left). The sharpening model predicts that BOLD activity in stimulus-non-selective (stimulus-selective) voxels decreases (stays unchanged) (right).

To test these predictions, the decoding accuracy as a function of moving windows would be generated according to the procedure outlined above. Linear regressions were fitted to the decoding accuracy time course and the slope was subjected to statistical analysis.

*Temporal dynamics of BOLD response in stimulus-selective and stimulus-non-selective voxels:* The fatigue and sharpening models make distinct predictions regarding the response dynamics of stimulus-selective and stimulus-non-selective voxels. According to the fatigue model, the BOLD activation in stimulus-selective voxels should exhibit decline, whereas the BOLD activation in stimulus-non-selective voxels should show minimal change (Figure 3B left). In contrast, the sharpening model predicts a decline in the BOLD activation in stimulus-non-selective voxels, whereas the BOLD activation in stimulus-selective voxels is expected to exhibit minimal change (Figure 3B right).

To test these predictions, for a given Gabor patch, the BOLD activation in voxels selective for the Gabor patch was first averaged for a trial, which was then averaged across subjects, the resulting value was then displayed as a function of trials. Similar temporal activation function of trials can be constructed for the stimulus-non-selective voxels. Linear regressions were fitted to the BOLD activation time courses of two types of voxels and the slope was subjected to statistical analysis.

## Results

fMRI data were recorded while subjects passively viewed the repeated presentation of two Gabor patches (Gabor 1=45° and Gabor 2=135°). Each Gabor was presented 60 times for a total of 120 trials. The order of presentation was randomized. We analyzed the fMRI data using both univariate and multivariate methods to assess learning and plasticity in the primary visual cortex (V1).

### Univariate analysis

For each Gabor 1 trial, the BOLD activation was averaged across voxels of V1 and then across all subjects. Repeating the procedure for all 60 repetitions resulted in the BOLD activation time course for Gabor 1. The BOLD activation time course for Gabor 2 was similarly obtained. The two time courses were then averaged and shown in Figure 4. Two types of behavior were observed. For the first 12 repetitions of the stimuli, labeled S1 (Stage1), the repetition suppression effect was apparent, whereas for the remaining repetitions, labeled S2 (Stage 2), no clear trend could be detected. For S1, linear regression models were fit to each subject’s data, and the slopes of regression models (-0.084 ± 0.048) were found to be significantly negative at p=0.027. Furthermore, the BOLD activation of the twelfth repetition was significantly lower than that of the first presentation at p<0.05. For S2, the slopes of the linear regression models were not significantly different from 0 (p=0.65) (Figure 4).

**Figure 4.**
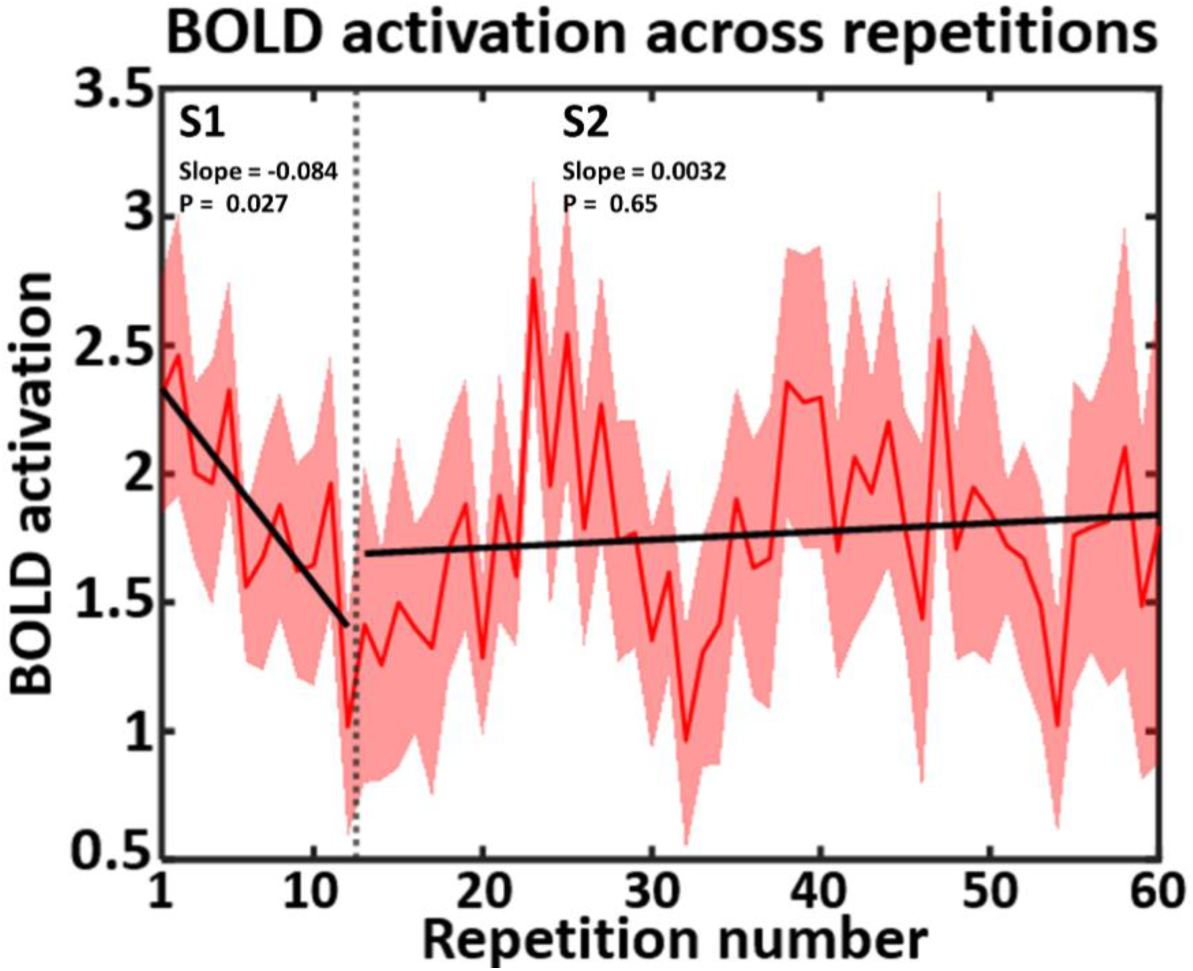
Univariate analysis of fMRI data. BOLD activation as a function of the repetition number was shown. Repetition suppression was observed for the first 12 repetitions of the stimuli (Stage 1 or S1). The remaining repetitions constituted Stage 2 or S2 during which significant fluctuations were seen with no systematic trend.

### Multivariate analysis: Stage 1

Having observed repetition suppression in S1, we next applied MVPA to examine the effect from the point of view of multivoxel patterns. Specifically, we decoded the differences between the multivoxel responses evoked by Gabor 1 and Gabor 2 using a moving widow approach, where the window length was 7 repetitions (each window was denoted by the repetition number of the middle trial) and the step size was 1 repetition. The decoding accuracy between the two Gabors was measured in each analysis window and averaged across 18 participants. The decoding accuracy time course in Figure 5A showed that the decoding accuracy in Window 1 was 59%, which is significantly higher than the chance level of 50% (p<0.01), increased to 73% (p<0.001) in Window 2, and then decreased to 54% at the end of S1, which was not significantly above chance (p=0.19). The generally decreasing trend of the decoding accuracy, which accorded with the generally decreasing trend of univariate BOLD activation in Figure 4 (S1), shows that at the multivoxel pattern level, Gabor 1 and Gabor 2 evoked less distinguishable patterns toward the end of repetition suppression. Regression fits to the data yielded a slope of -2.09 ± 1.07 which was significantly less than 0 at p<0.05 (Figure 5A). The diminishing decoding accuracy, according to the model formulation in multivoxel terms in Figure 3A, is consistent with the prediction of the fatigue model.

**Figure 5.**
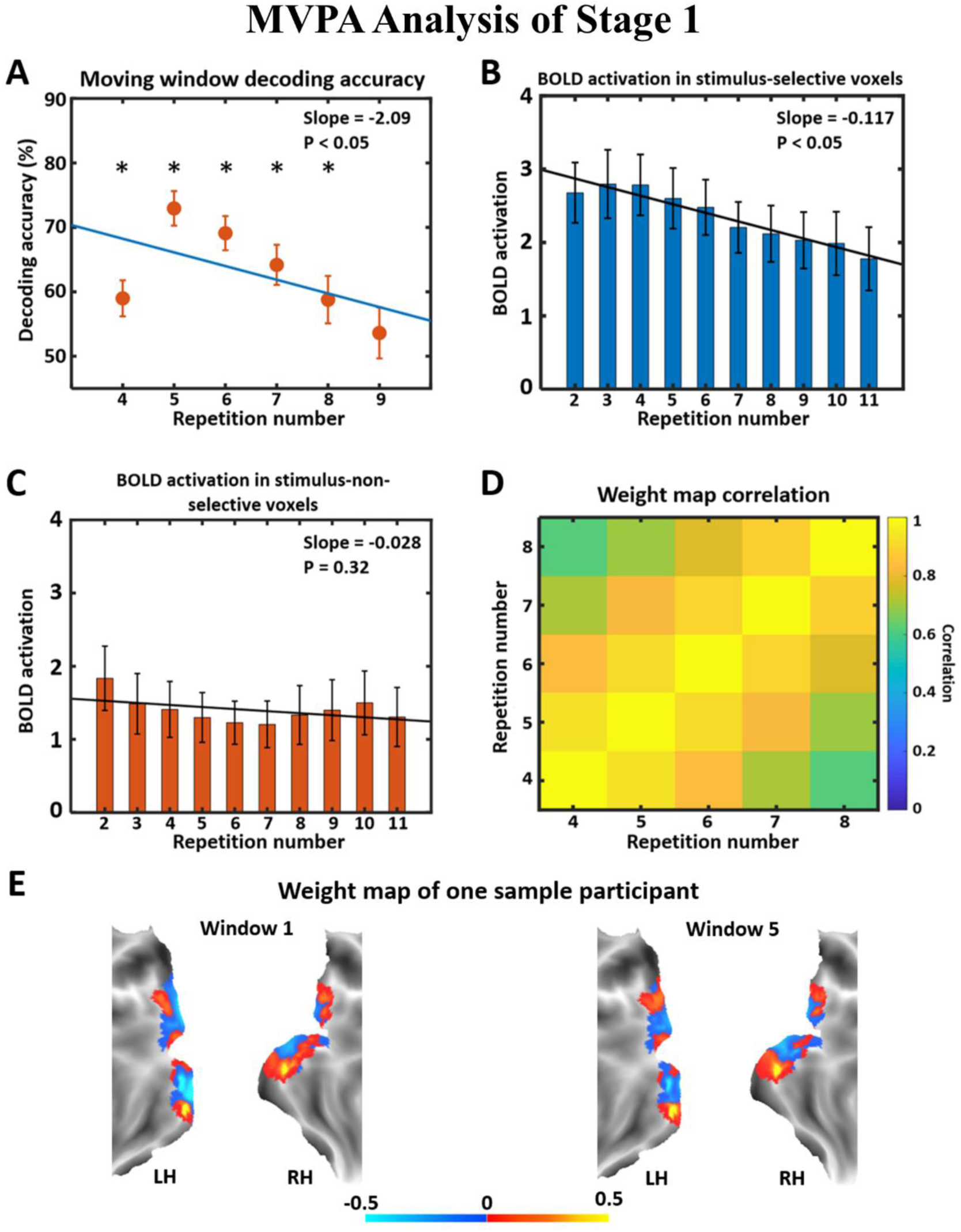
MVPA analysis of Stage 1 learning dynamics. **(A)** Decoding accuracy in V1 across stimulus repetition. The asterisk indicates decoding accuracy significantly above chance level (50%) at p<0.05. The repetition number is the trial number in the middle of each window. **(B)** BOLD activation time course in stimulus-selective voxels. **(C)** BOLD activation time course in stimulus-non-selective voxels. **(D)** Weight map correlation for all combinations of moving windows. Only weight maps in decodable windows were used. **(E)** Weight maps from Window 1 and Window 5 of one sample participant are shown on the flattened brain.

To test this idea further, we utilized the weight map analysis and defined two types of voxels: one that selectively responds to Gabor 1 and one that selectively responds to Gabor 2. For stimulus-selective voxels, responses of Gabor 1-selective voxels to Gabor 1 and responses of Gabor 2-selective voxels to Gabor 2 were averaged and plotted as a function of stimulus repetition. As shown in Figure 5B, the responses of the stimulus-selective voxels exhibited a statistically declining trend, with the slope being -0.117 ± 0.06 which is significantly less than 0 at p<0.05. For the stimulus-non-selective voxels, responses of Gabor 1-selective voxels to Gabor 2 and responses of Gabor 2-selective voxels to Gabor 1 were averaged and plotted as a function of stimulus repetition. The slope was - 0.028 ± 0.06 which was not significantly different from 0 (p=0.32) (Figure 5C). Based on Figure 3B, these weight map results demonstrated that the fatigue model accounted for the neural dynamics underlying Stage 1 learning, consistent with the conclusion drawn from the decoding accuracy dynamics.

Whether the neural representations of the two stimuli were stable during the course of repetition suppression was tested by computing correlation across weight maps for all pairwise combinations of moving windows. As shown in Figure 5D, all the correlations were significantly positive (p<0.05), demonstrating that the patterns underlying the two stimuli stayed relatively stable. Figure 5E shows the weight maps from Window 1 and Window 5 for a typical subject. The consistency between the two maps was readily noticeable.

### Multivariate analysis: Stage 2

As seen in Figure 4, following the initial repetition suppression, the univariate analysis revealed fluctuating BOLD activity with no systematic trend. We then applied moving-window MVPA to data from this stage of the task, referred to as Stage 2 (S2). As shown in Figure 6A, at the beginning of S2, the decoding accuracy was at chance level, but with further repetitions of the stimuli, the decoding accuracy begins to trend up, reaching statistical significance at Window 4 (R25 – R44) and staying above chance level until the end of the session. Linear regression fitting of the data yielded a slope of 1.94 ± 0.26 which was significantly larger than 0 at p<0.0001 (Figure 6A). Thus, beyond the initial repetition suppression, there was a second-stage of perceptual learning, in which the neural representations of the two Gabor patches become more distinct from one another with learning, in contrast to what was observed during repetition suppression learning (Stage 1) where the neural representations of the two Gabors became less distinct with learning. According to the multivoxel formulation of the models in Figure 3A, the above temporal behavior of the decoding accuracy is consistent with a sharpening process.

**Figure 6.**
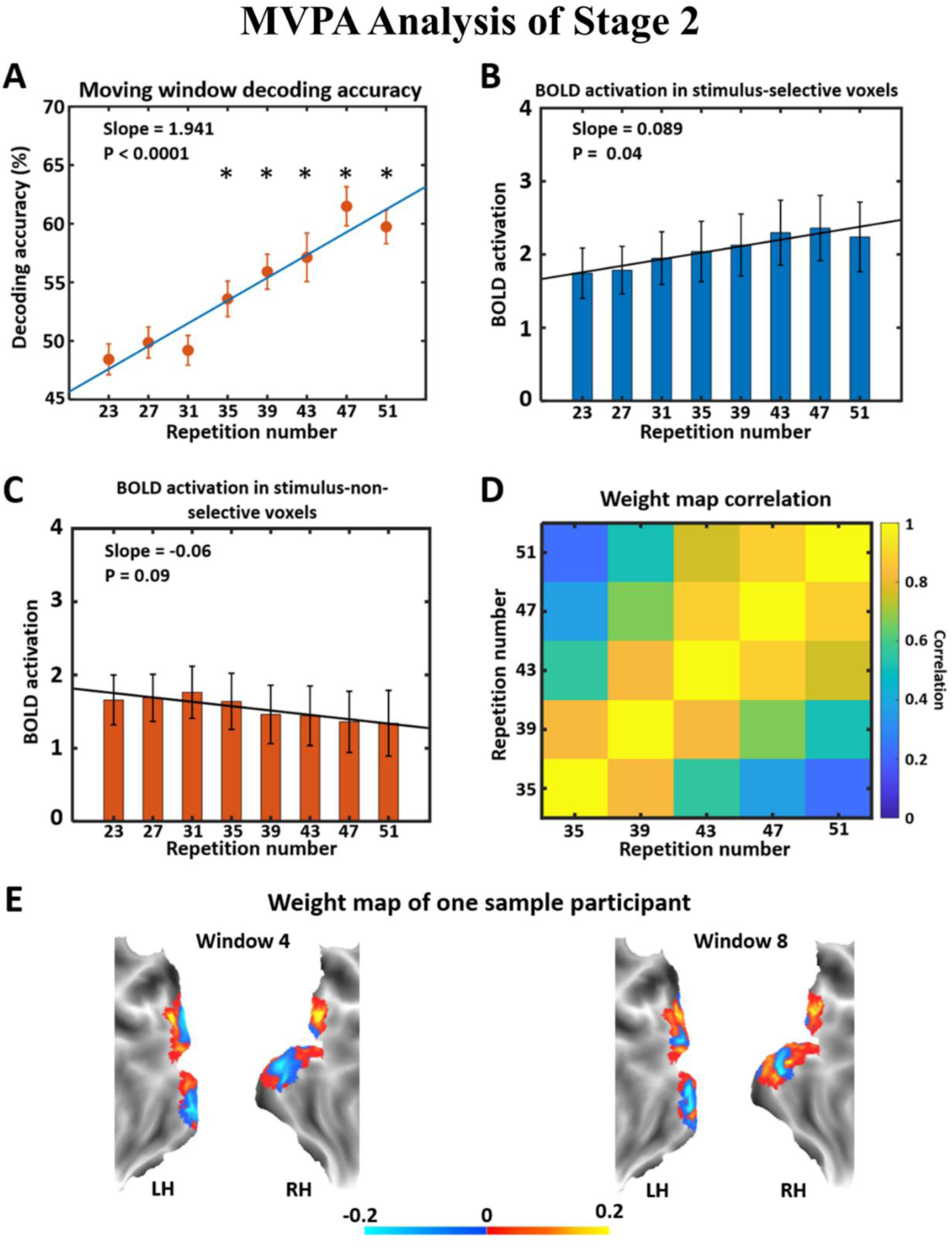
MVPA analysis of Stage 2 learning. **(A)** Decoding accuracy in V1 across stimulus repetition. The asterisk indicates decoding accuracy significantly above chance level (50%) at p<0.05. The repetition number denotes the trial number in the middle of each window. **(B)** BOLD activation time course in stimulus- selective voxels. **(C)** BOLD activation time course in stimulus-non-selective voxels. **(D)** Weight map correlation for all combinations of moving windows. Only weight maps in decodable windows were used. **(E)** Weight maps from Window 4 and Window 8 of one sample participant are shown on the flattened brain.

We examined this idea further using the weight map analysis. For stimulus-selective voxels, responses of Gabor 1-selective voxels to Gabor 1 and responses of Gabor 2-selective voxels to Gabor 2 were averaged and plotted as a function of stimulus repetition. As shown in Figure 6B, the responses exhibited an increasing trend, where the slope was 0.089 ± 0.05 which was statistically significantly larger than 0 at p=0.04. For stimulus-non-selective voxels, responses of Gabor 1-selective voxels to Gabor 2 and responses of Gabor 2-selective voxels to Gabor 1 were averaged and plotted as a function of stimulus repetition. A declining trend was seen although the slope was only marginally statistically significant (p=0.09) (Figure 6C). Thus, the weight map analysis results are in line with the earlier conclusion from decoding accuracy analysis, suggesting that the sharpening model accounted for the neural dynamics underlying Stage 2 learning. It is worth noting that the sharpening process seen here is slightly different from the one depicted in Figure 3B, in that the activity in the stimulus-selective voxels exhibited significant increase rather than remaining relatively constant.

The weight map correlation analysis in Figure 6D shows that the neural representations generally stayed stable during the course of S2. All the correlations, except the one for analysis windows centered on repetition 35 and 51, were significantly positive (p<0.05), demonstrating that the patterns underlying the two stimuli stayed relatively stable. Figure 6E shows the weight maps from Window 4 and Window 8 for a typical subject.

### Multivariate analysis: Comparison between Stage 1 and Stage 2

At the beginning of Stage 1 and during much of Stage 2, the two Gabor patches are decodable based on their distinct activation patterns in V1. The question is whether the decodable representations of Gabor 1 and 2 are shared across the two stages of learning. If the representations of Gabor 1 and Gabor 2 are similar between the two stages, then MVPA classifiers constructed on data from one stage would decode the data from another. But as shown in Figure 7A, the cross-decoding accuracy was at chance level for all possible window combinations, indicating that during Stage 2 learning, new representations for the two Gabor patches evolved, which were different from that during Stage 1. The weight map correlation analysis further supports this observation, showing that the weight maps from S1 and S2 are not correlated for all possible window combinations (Figure 7B), in contrast to the within-stage weight map correlation in Figures 5D and 6D. To further visualize this, we plotted the weight maps from S1 and S2 for two sample participants. In these weight maps, the cooler color (blue) represents voxels selective for Gabor 1 and the warmer color (yellow) Gabor 2. Visually, it is apparent that voxels that are selective for Gabor 1/Gabor2 in Stage 1 differ from those selective for Gabor 1/Gabor 2 in Stage 2 (Figure 7C), which accorded with the chance-level cross-decoding result and the zero correlation result between weight maps above. Furthermore, as can be observed in Figure 7C, the dissimilarity in neural patterns is not only evident between learning stages within the same participant but is also evidence between different participants within the same learning stage, suggesting that even for simple stimuli such as Gabor patches and in early visual cortex such as V1, the neural representations are individual-specific.

**Figure 7.**
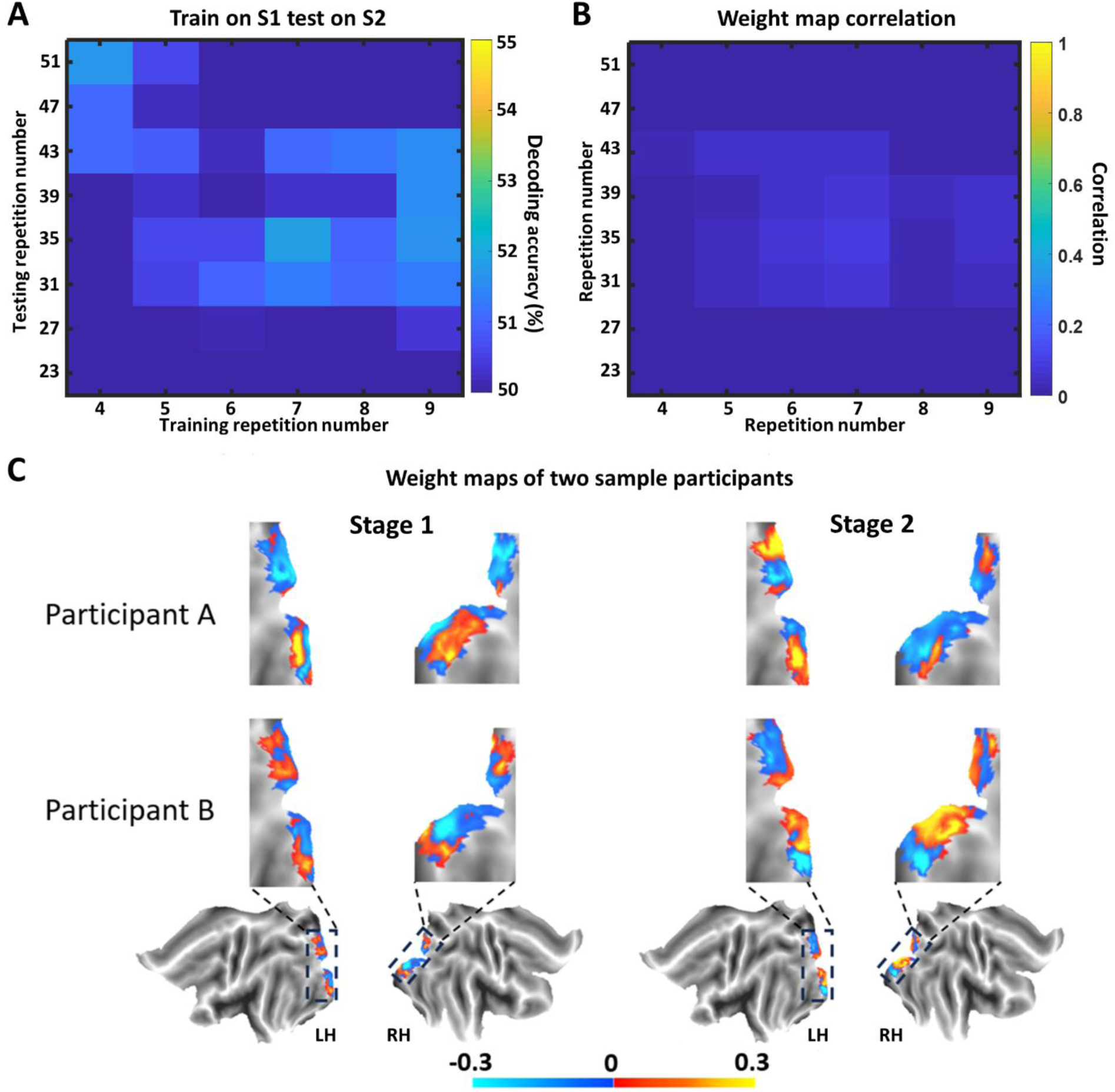
Cross-stage decoding and weight map analysis. **(A)** Classifiers trained on analysis windows in Stage 1 were applied to test data from Stage 2. The cross-decoding accuracy was at chance level (50%). The repetition number indicates the middle trial in each window. **(B)** Weight map correlations for all cross-stage combinations of analysis windows were not significantly different from 0. **(C)** Weight maps from two participants in V1 are shown on the flattened hemisphere. The warmer color (yellow) indicates voxels that are selective for Gabor 1 and the cooler color (blue) indicates voxels that are selective for Gabor 2.

## Discussion

We investigated the effects of stimulus repetition on the neural representations of the repeating stimuli in primary visual cortex. Human observers viewed 120 presentations of two Gabor patches (each repeating 60 times) in random order while their fMRI was recorded. Univariate analysis revealed two types of learning behavior in V1: (1) BOLD activation decreased over the first twelve repetitions of the stimuli (referred to as Stage 1), replicating the well-known effect of repetition suppression and (2) BOLD activation for the remaining 48 repetitions of the two stimuli, denoted Stage 2, exhibited significant fluctuations but no systematic trend. Applying MVPA decoding using a moving window approach to Stage 1, we found that (1) the decoding accuracy between the two Gabors went from being above chance level (∼60% to ∼70%) at the beginning of the stage to being at chance level at the end of the stage (∼50%). This result, along with the attendant weight map analysis, suggested that the learning dynamics in Stage 1 were consistent with the predictions of the fatigue model. For Stage 2, the accuracy of decoding between the two Gabor patches was at chance level initially and became progressively higher as stimulus repetition continued, rising above chance level at ∼35th repetition and staying above chance level until the end of the experiment. This result, along with the attendant weight map analysis, suggests that a sharpening process underlies the learning dynamics in Stage 2. A cross-decoding analysis shows that the two Gabor patches are represented differently in Stage 1 vs Stage 2, indicating that the same physical stimuli evoked distinct multivoxel patterns depending on the stage of learning.

### Stage 1 learning: Repetition fatigue

Repetition suppression is a robust index of short-term perceptual learning and has been demonstrated extensively in both humans and animals (Sobotka et al. 1996; Naccache et al. 2001). For example, a single-unit study in nonhuman primates has reported that neurons in IT show a continual reduction in firing rate with each stimulus presentation for up to six to ten repetitions, after which the response became stable (Li et al. 1993; Sawamura et al. 2006). fMRI studies have found that the first few stimulus repetitions produced a substantial reduction in signal strength in LOC and no further reduction was observed upon continued repetitions of the stimuli (Grill-Spector et al. 2001; Sayres et al. 2006). Our results replicate these findings in V1. Specifically, averaging BOLD activation across all V1 voxels for Gabor 1 and for Gabor 2 and plotting the result as a function of repetition, we observed repetition suppression for the first twelve repetitions of the stimuli, after which the univariate BOLD response showed no systematic trend.

Among the models that have been proposed to account for the neural mechanisms of repetition suppression, the fatigue model posits that stimulus repetition reduces the responsiveness of neurons selective for the repeating stimuli (Grill-Spector et al. 2006; Carandini et al. 1997; Carandini 2000), whereas the sharpening model, in contrast, posits that stimulus repetition sharpens neuronal tuning by reducing the activity of neurons not selective for the repeating stimuli; both models can explain the reduction of responses at the population level (Grill-Spector et al. 2006; Wiggs et al. 1998). Evidence in support of one or the other model has appeared in single-unit studies. For example, Freedman et al. and Baker et al. both demonstrated that the repetition of stimuli leads to increased selectivity in the neural responses with a concurrent attenuation of activity in neurons that are less selective (Freedman et al. 2006; Baker et al. 2002), aligning with the predictions of the sharpening model. Conversely, Li et al. reported that the most significant reduction in neuronal response occurs for those stimuli that initially elicited the largest response, a finding that lends support to the fatigue model (Li et al. 1993).

In humans, the testing of these models has proven to be not straightforward. Despite that repetition suppression has been consistently observed with fMRI in a variety of brain regions, ranging from V1 to higher order visual regions such as IT to executive structures in the frontal lobe (Weigelt et al. 2008; Buckner et al. 1998; Kourtzi et al. 2001; Sawamura et al. 2005; Ewbank et al. 2005; Pourtois et al. 2009), the commonly applied univariate fMRI approach, which involves the averaging of activities across voxels in a given region of interest, lacks the requisite spatial resolution and response specificity for testing the models of repetition suppression. Weiner et al. examined neural responses of stimulus-selective voxels under novel and repeated stimulus conditions, revealing a reduction in the fMRI signal for repeated stimuli (Weiner et al. 2010). Their study did not consider activities in stimulus-non-selective voxels. The lack of single-trial data limited the investigation of neural representation dynamics.

Over the past two decades, the advent of the MVPA method has opened new avenues for addressing challenging neuroscience questions, yielding insights not possible with the univariate approach (Said et al. 2010; Bo et al. 2021; Visser et al. 2013; Čeko et al. 2022; Bo et al. 2022). In a typical MVPA analysis, two experimental conditions are compared by training a SVM classifier on the neural data evoked by the two experimental conditions, with above-chance classification or decoding accuracy taken as the evidence that the two experimental conditions evoke distinct neural patterns and the brain region is involved in the cognitive operation under investigation. For the present study, the two experimental conditions are the two Gabor patches that are both repeating. By employing a moving window approach and performing MVPA decoding within each window, we examined how decoding accuracy evolves as a function of repetition. For the fatigue model to hold, the decoding accuracy between the two Gabors should decrease with stimulus repetition, whereas for the sharpening model to hold, the decoding accuracy between the two Gabors should increase with stimulus repetition. The result from the MVPA analysis, demonstrating decreasing decoding accuracy, suggests that the repetition suppression observed during Stage 1 results from a fatigue process. A similar finding has been made in the lateral ventral temporal cortex using complex visual stimuli (Weiner et al. 2010).

The weight map is another tool that can be derived from the SVM classifier. In particular, the transformation proposed by Haufe et al. (Haufe et al. 2014) enables the dividing of voxels in a ROI into those that are selective for Gabor 1 and for Gabor 2, respectively. By assessing the neural response in voxels that are selective for a given stimulus (stimulus-selective voxels) and the neural responses in voxels that are not selective for a given stimulus (stimulus-non-selective voxels), we are then equipped with the ability to test the models of repetition suppression in a way that is analogous to the single unit studies in nonhuman primates. For the fatigue model to hold, neural activity in the stimulus-selective voxels should decrease and that in the stimulus-non-selective voxels should remain steady, whereas for the sharpening model to hold, neural activity in the stimulus-selective voxels should remain steady and that in the stimulus-non-selective voxels should decrease (Figure 3B). The result from the weight map analysis, demonstrating decreasing neural activity in stimulus-selective voxel and steady neural activity in stimulus-non-selective voxels, supports the conclusion above that repetition suppression in Stage 1 results from a fatigue process.

### Stage 2 learning: Repetition sharpening

As indicated earlier, repetition suppression reflects neural plasticity over the short term, which in our data corresponds to ∼10 repetitions of each stimulus (Stage 1). Over the next ∼50 repetitions of each of the two Gabors, referred to as Stage 2, univariate BOLD activity from V1 exhibited significant fluctuations but no systematic trend. Suspecting that the univariate analysis lacks the sensitivity to uncover the underlying learning effects (de Beeck et al. 2006), we applied the MVPA approach to complement the univariate analysis. The MVPA results indicated that the neural response patterns to Gabor 1 and Gabor 2 continued to evolve, becoming more distinct from one another, as evidenced by increasing decoding accuracy. From the weight map analysis, the BOLD response in stimulus-selective voxels increased with learning, whereas the BOLD response in stimulus-non-selective voxels decreased with learning, which agrees with the conclusion that a sharpening process underlies the second stage of repetition learning.

Repeated stimulus exposure is often linked to improvement in stimulus discrimination and memory. The repetition suppression observed in Stage 1 is unlikely the reason for such improvement due to the fact that the distinctiveness of the two stimulus-evoked neural patterns decreased with learning. Behavioral studies often find that detectable improvement in discriminative ability requires large amounts of repeated exposures to stimuli. For instance, perceptual performance improvement occurs after thousands of repetitions across days in a variety of perceptual tasks, including luminance contrast detection (Sowden et al. 2002), orientation detection (Censor et al. 2006, Zhang et al. 2020), object recognition (Furmanski et al. 2000; de Beeck et al. 2006) and color discrimination (Horiuchi et al. 2024). Despite substantial behavioral evidence, there remains a dearth of research on the neural underpinnings behind improved behavior during longer periods of task-free learning. One prior study found that memory performance improvements are associated with increased dissimilarity in neural patterns in the hippocampal region (LaRocque et al. 2013). Since increased dissimilarity in neural patterns is equivalent to increased decoding accuracy, our study can be seen as providing sharpened neural representations as a possible mechanism for previously observed improvement in perceptual abilities through extensive stimulus exposure.

### Stage-dependent neural representations of Gabor 1 and Gabor 2

In both Stage 1 and Stage 2, the two Gabors were decodable in part of the stage, i.e., the early part of Stage 1 and the late part of Stage 2. An obvious question is whether the neural representations of Gabor 1 and Gabor 2 are the same across the two stages. Previous work has shown that neural representations drift with learning both in single-unit recordings (LeMessurier et al. 2018; Carandini 2004), where the timescale over which this occurs ranges from minutes to weeks (Lütcke et al. 2013; Deitch et al. 2021; Xia et al. 2021), and in human fMRI research (Roth et al. 2023). To address this question, we conducted a cross-stage decoding analysis: classifiers were trained on the data from the Stage 1 and tested on the data from the Stage 2. Chance level cross decoding accuracy (∼50%) was found. This cross-decoding finding was further supported by cross-stage weight map correlations which were found to be not different from 0. These results, suggesting that Gabor 1 and Gabor 2 evoked different patterns of neural activity in Stage 1 vs Stage 2, is in contrast to the within-stage weight map correlation analysis showing that the patterns of neural activity evoked by Gabor 1 and Gabor 2 remained relatively stable within each stage of learning.

### Methodological considerations

The findings reported here are made possible by several features of our methodology. First, in our study, we contrasted two stimuli that were both repeating. Previous research has typically compared repeating stimuli with novel stimuli. It has been pointed out that some of the observed repetition effects may be attributable to the surprise effects of the novel stimuli (Schomaker et al. 2015). This concern does not apply to our experiment. In addition, using the two-repeating-stimuli paradigm, we were able to reformulate the models of repetition suppression in terms of multivariate representations, enabling the testing of the repetition suppression models in fMRI data. Second, to evaluate the hypothesis regarding models of repetition suppression at a multivariate level, decoding accuracy alone is insufficient for a comprehensive evaluation of different models, as identical decoding dynamics can result from different underlying voxel dynamics. For example, an increase in decoding accuracy could be driven either by an increase in BOLD activation in stimulus-selective voxels while activation in stimulus-non-selective voxels remains unchanged, or by stable activation in stimulus-selective voxels while activation in stimulus-non-selective voxels decreases. The weight map analysis provides the necessary insights to disambiguate and thus complements the decoding accuracy dynamics. Third, the MVPA results during Stage 2, while suggesting a sharpening process, are not strictly in accordance with the definition of sharpening in the literature: both enhancement of activation in stimulus-selective voxels and suppression of activation in stimulus-non-selective voxels were observed in our data whereas the traditional definition of sharpening places emphasis on the decreased activity in stimulus-non-selective neurons. The weight map analysis is again essential in establishing this finding.

## Summary and outlook

In this study we reported the discovery of a two-stage perceptual learning process in primary visual cortex during passive viewing of a large number of repetitions of two Gabor patches. The first stage of learning is characterized by repetition suppression undergirded by a fatigue process whereas the second stage of learning is characterized by the graduate emergence of distinct neural representations of the two Gabors undergirded by a sharpening process. Two factors played a key role in enabling these findings: (1) the substantial number of repetitions of two stimuli allowed learning processes to fully unfold and (2) the MVPA decoding method and the associated weight map analysis made it possible to characterize the hypothetical neural mechanisms underlying these learning processes. A major limitation of the present study is that there was no behavioral testing to accompany the neural recordings. Thus, the behavioral consequences of the main findings remain to be assessed in future studies.

## Acknowledgements

This work was supported by NIH grant MH125615.

